# Genome-wide association meta-analysis for early age-related macular degeneration highlights novel loci and insights for advanced disease

**DOI:** 10.1101/2019.12.20.883801

**Authors:** Thomas W Winkler, Felix Grassmann, Caroline Brandl, Christina Kiel, Felix Günther, Tobias Strunz, Lorraine Weidner, Martina E Zimmermann, Christina A. Korb, Alicia Poplawski, Alexander K Schuster, Martina Müller-Nurasyid, Annette Peters, Franziska G Rauscher, Tobias Elze, Katrin Horn, Markus Scholz, Marisa Cañadas-Garre, Amy Jayne McKnight, Nicola Quinn, Ruth E Hogg, Helmut Küchenhoff, Iris M Heid, Klaus J Stark, Bernhard HF Weber

## Abstract

**Background:** Advanced age-related macular degeneration (AMD) is a leading cause of blindness. While around half of the genetic contribution to advanced AMD has been uncovered, little is known about the genetic architecture of the preceding early stages of the diseases.

**Methods:** To identify genetic factors for early AMD, we conducted a genome-wide association meta-analysis with 14,034 early AMD cases and 91,214 controls from 11 sources of data including data from the International AMD Genomics Consortium (IAMDGC) and the UK Biobank (UKBB). We ascertained early AMD via color fundus photographs by manual grading for 10 sources and by using an automated machine learning approach for >170,000 images from UKBB. We searched for significant genetic loci in a genome-wide association screen (P<5×10^-8^) based on the meta-analysis of the 11 sources and via a candidate approach based on 13 suggestive early AMD variants from Holliday et al 2013 (P<0.05/13, additional 3,432 early AMD cases and 11,235 controls). For the novel AMD regions, we conducted in-silico follow-up analysis to prioritize causal genes and pathway analyses.

**Results:** We identified 11 loci for early AMD, 9 novel and 2 known for early AMD. Most of these 11 loci overlapped with known advanced AMD loci (near *ARMS2/HTRA1, CFH, APOE, C2, C3, CETP, PVRL2, TNFRSF10A, VEGFA*), except two that were completely novel to any AMD. Among the 17 genes within the two novel loci, in-silico functional annotation suggested *CD46* and *TYR* as the most likely responsible genes. We found the presence or absence of an early AMD effect to distinguish known pathways of advanced AMD genetics (complement/lipid pathways or extracellular matrix metabolism, respectively).

**Conclusions:** Our data on early AMD genetics provides a resource comparable to the existing data on advanced AMD genetics, which enables a joint view. Our large GWAS on early AMD identified novel loci, highlighted shared and distinct genetics between early and advanced AMD and provides insights into AMD etiology. The ability of early AMD effects to differentiate the major pathways for advanced AMD underscores the biological relevance of a joint view on early and advanced AMD genetics.

## BACKGROUND

Age-related macular degeneration (AMD) is the leading cause of irreversible central vision impairment in industrialized countries. Advanced AMD presents as geographic atrophy (GA) and/or neovascular (NV) complications (1). Typically, advanced AMD is preceded by clinically asymptomatic and thus often unrecognized early disease stages. Early AMD is characterised by differently sized yellowish accumulations of extracellular material between Bruch’s membrane and retinal pigment epithelium (RPE) or between RPE and the photoreceptors (drusen or subretinal drusenoid deposits, respectively). Other features of early AMD are RPE abnormalities, including depigmentation or increased amount of pigment (1).

Early and advanced AMD can be documented by color fundus imaging of the central retina and/or other multimodal imaging approaches including optical coherence tomography (OCT) (1–3). While the definition of advanced AMD is reasonably homogeneous across clinical and epidemiological studies, the classification of early AMD is more variable and different studies traditionally apply differing classification systems (4,5).

Epidemiological studies show that high age is the strongest risk factor for early and advanced AMD onset as well as progression (1,6–8). A robust genetic influence was shown for advanced AMD (1,9,10) with 34 distinct loci at genome-wide significance in a large genome-wide association study (GWAS) for advanced AMD (9). The genes underneath these advanced AMD loci were found to be enriched for genes in the alternative complement pathway, HDL transport, and extracellular matrix organization and assembly (9).

Exploring the genetics of early AMD offers the potential to understand the mechanisms of early disease processes, but also for the development to advanced AMD when comparing genetic effect sizes for early and advanced stages. Yet there have been few published GWAS searches for early AMD. One meta-analysis on 4,089 early AMD patients and 20,453 control persons reported two loci with genome-wide significance, both being well known from advanced AMD, the *CFH* and the *ARMS2/HTRA1* locus (11).

We have thus set out to gather GWAS data for early AMD from 11 sources including own study data, data from the International AMD Genomics Consortium (IAMDGC), dbGaP and UK Biobank to conduct the largest GWAS meta-analysis on early AMD to date.

## METHODS

### GWAS data from 11 sources

We included 11 sources of data with GWAS data and color fundus photography for early AMD phenotyping (**Table S1**). Our studies were primarily population-based cohort studies, where the baseline survey data were used for this analysis from studies of the authors (GHS, LIFE, NICOLA, KORA, AugUR) as well as for publicly available studies from dbGaP (ARIC, CHS, WHI; accession numbers: phs000090.v5.p1, phs000287.v6.p1, phs000746.v2.p3). We also included data from UK Biobank for participants from baseline and additional participants from the follow-up survey, since the color fundus photography program had started only after the main study onset (application number #33999). The studies captured an age range from 25 to 100 years of age (mean age from 47.5 years to 77.2 years across the 10 population-based studies, AugUR with the very old individuals range from 70.3 years to 95 years). About 50% of the study participants in each study were male (except for the Women’s Health Initiative, WHI) and all demonstrated European ancestry. For each of these cross-sectional data sets, participants with at least one eye gradable for AMD (see below) and with existing GWAS data were eligible for our analysis. We excluded participants with advanced AMD. We used participants with ascertained early AMD as cases and participants being ascertained for not having any signs of AMD as controls (n=7,363 cases, 73,358 controls across these population-based studies). Case-control data were also included from IAMDGC (http://amdgenetics.org/). The early AMD GWAS from IAMDGC is based on 24,527 individual participant data from 26 sources (9). This data includes 17,856 participants with no AMD and 6,671 participants with early AMD (excluding the 16,144 participants with advanced AMD). The cases and controls from IAMDGC were 16 to 102 years of age (mean age = 71.7y). For all of these participants, DNA samples had been gathered and genotyped centrally (see below) (9).

### Genotyping and Imputation

All population-based studies were genotyped, quality controlled and imputed using similar chip platforms and imputation approaches (**Table S2**). As the imputation backbone, the 1000 Genomes Phase 1 or Phase 3 reference panel was applied (12), except GHS was imputed based on the Haplotype Reference Consortium (HRC) (13) and UK Biobank was imputed based on HRC and the UK10K haplotype resource (14). Details on the UK Biobank genotypic resource are described elsewhere (15). For the IAMDGC case-control data, DNA samples had been gathered across all participants and genotyped on an Illumina HumanCoreExome array and quality controlled centrally. Genotype quality control and imputation to the 1000 Genomes phase 1 version 3 reference panel (>12 million variants) were conducted centrally. Details on the IAMDGC data were described in detail by Fritsche et al (9).

### Phenotyping

Across all studies included into this analysis, early AMD and the unaffected status was ascertained by color fundus photography. For participants from AugUR and LIFE, “early AMD” was classified according to the Three Continent Consortium (3CC) Severity Scale (4), which separates “mild early” from “moderate” and “severe early” AMD stages depending on drusen size, drusen area, or the presence of pigmentary abnormalities (4). For the analysis, we collapsed any of these “early” AMD stages into the definition of “early AMD”. However, the 3CC Severity Scale was not available for the other studies. In these, similar early AMD classifications, considering drusen size or area and presence of pigmentary abnormalities, were used (**Table S1**): For participants from GHS, the Rotterdam Eye Study classification was applied (16). For participants from NICOLA, the Beckman Clinical Classification was utilized (17). Participants from the KORA study were classified as “early AMD” based on the AREDS-9 step classification scheme and we defined “early AMD” for this analysis by AREDS-9 steps 2-8 (18). The ascertainment of IAMDGC study participants is described in detail elsewhere and covers various classification systems (9). Of note, LIFE and NICOLA phenotyping incorporated OCT information additional to the information from color fundus imaging (**Table S1**). For UK Biobank participants, color fundus images were received (application number 33999); there was no existing AMD classification available in UK Biobank (see below). The AMD status of a person was derived based on the AMD status of the eye with the more severe AMD stage (“worse eye”) when both eyes were gradable, and as the grade of the one available eye otherwise. Eyes were regarded as gradable, if at least one image of the eye fulfilled defined quality criteria allowing for the assessment of AMD (bright image, good color contrast, full macular region captured on images). Images were excluded from AMD grading if they revealed obscuring lesions (e.g. cataract) or lesions considered to be the result of a competing retinal disease (such as advanced diabetic retinopathy, high myopia, trauma, congenital diseases, or photocoagulation unrelated to choroidal neovascularization). Details for IAMDGC are described previously (9). Persons with gradable images for at least one eye were included in this analysis. Persons with advanced AMD defined as presence of neovascularization or geographic atrophy in at least one eye were excluded for the main GWAS on early AMD.

### Automated classification of early AMD in UK Biobank

To obtain early AMD phenotype data for UK Biobank participants, we used a pre-trained algorithm for automated AMD classification based on an ensemble of convolutional neural networks (19). In the UKBB baseline data, fundus images were available for 135,500 eyes of 68,400 individuals with at least one image. Among the additional 38,712 images of 19,501 individuals in the follow-up, there were 17,198 individuals without any image from baseline. For each image (eye) at baseline and follow-up, we predicted the AMD stage on the AREDS-9 step severity scale using the automated AMD classification. We defined a person-specific AMD stage at baseline and follow-up based on the worse eye. Eyes that were classified as ungradable were treated as missing data and, if diagnosis was available for only one eye, the person-specific AMD stage was based on the classification of the single eye. If we obtained an automated disease classification to an AMD stage (i.e. not “ungradable” for both eyes) at baseline and follow-up, we used the follow-up disease stage (and follow-up age) in the association analysis. By this, we obtained an automated AMD classification for 70,349 individuals (2,161 advanced AMD, 3,835 early AMD, 64,353 unaffected). Individuals with advanced AMD were excluded from this analysis. Finally, we yielded 57,802 unrelated individuals of European ancestry with valid GWAS data that had either early AMD or were free of any AMD (3,105 cases, 54,697 controls). We evaluated the performance of the automated disease classification by selecting 2,013 individuals (4,026 fundus images) and manual classification based on the 3CC Severity Scale. We found substantial agreement between the automated and the manual classification for the four categories of “no AMD”, “early AMD”, “advanced AMD” and “ungradable” (concordance=79.5%, Cohen’s kappa κ=0.613) (20).

### Study-specific association analyses

Study-specific logistic regression analyses (early AMD cases versus controls, excluding advanced AMD cases) were applied by study partners (in Regensburg, Leipzig, Mainz, Belfast) using an additive genotype model and according to a pre-defined analysis plan. All publicly available data from dbGAP (studies ARIC, CHS and WHI) and UK Biobank as well as IAMDGC data was analyzed in Regensburg. All studies inferred the association of each genetic variant with early AMD using a Wald test statistic as implemented in RVTESTS (21). Age and two principal components (to adjust for population stratification) were included as covariates in the regression models. The IAMDGC analyses were further adjusted for DNA source as done previously (9). For the IAMDGC data that stemmed from 26 sources, we conducted a sensitivity analysis additionally adjusting for source membership according to previous work highlighting slight differences in effect estimates (22); we found the same results.

### Quality control of study-specific aggregated data

GWAS summary statistics for all data sources were processed through a standardized quality-control (QC) pipeline (23). This involved QC checks on file completeness, range of test statistics, allele frequencies, population stratification as well as filtering on low quality data. We excluded variants with low minor allele count (MAC<10, calculated as MAC=2*N_eff_*MAF, with N_eff_ being the effective sample size, N_eff_=4N_Cases_*N_Controls_/(N_Cases_+N_Controls_) and MAF being the minor allele frequency), low imputation quality (rsq<0.4) or large standard error of the estimated genetic effect (SE>10). Genomic control (GC) correction was applied to each GWAS result to correct for population stratification within each study (24). The estimation of the GC inflation factor was based on variants outside of the 34 known advanced AMD regions (excluding all variants within <5 Mb base positions to any of the 34 known advanced AMD lead variants). The GC factors ranged from 1.00 to 1.04 (**Table S2**). We transferred all variant identifiers to unique variant names consisting of chromosomal, base position (hg19) and allele codes in (e.g. “3:12345:A_C”, allele codes in ASCII ascending order).

### Meta-analysis

For signal detection and effect quantification, study-specific genetic effects were combined using an inverse-variance weighted fixed effect meta-analysis method as implemented in METAL (25). We performed additional quality control on meta-analysis results: We only included variants for identification that were available (i) in at least two of the data sources with a total effective sample size of more than 5,000 individuals (N_eff_>5,000) and (ii) for chromosome and position annotation in dbSNP (hg19). A conservative second GC correction (again focusing on variants outside the known advanced AMD regions) was applied to the meta-analysis result, in order to correct for potential population stratification across studies (24). The GC lambda factor of the meta-analysis was 1.01. We judged the variants’ association at genome-wide significance level (P< 5×10^-8^). To evaluate the robustness of any novel genome-wide significant AMD locus, we performed leave-one-out (LOO) meta-analyses.

### Variant selection, locus definition, and independent replication

We combined genome-wide significant variants (P<5.0×10^-8^) into independent loci by using a locus definition similar to what was done previously (9): the most significant variant was selected genome-wide, all variants were extracted that were correlated with this lead variant (r^2^>0.5, using IAMDGC controls as reference) and a further 500 kb were added to both sides. All variants overlapping the so-defined locus were assigned to the respective locus. We repeated the procedure until no further genome-wide significant variants were detected. Genes overlapping the so-defined loci were used for biological follow-up analyses (gene region defined from start to end). To identify independent secondary signals at any novel AMD locus, approximate conditional analyses were conducted based on meta-analysis summary statistics using GCTA (26).

For an independent replication stage for genome-wide significant lead variants identified in our GWAS meta-analysis, we utilized the data from Holliday et al. (11) where possible. Since Holliday et al. did not provide genome-wide summary statistics, this was limited to variants, for which (or for reasonable proxies) summary statistics were reported. To account for the overlap between the current meta-analysis and the Holliday et al. meta-analysis, we removed the effects of the two overlapping studies (ARIC and CHS) from the reported Holliday et al. summary statistics. For β_Holl_, β_ARIC,_ β_CHS_ and SE_Holl_, SE_ARIC,_ SE_CHS_ being the genetic effects and standard errors of the Holliday et al. meta-analysis (reported) and of the ARIC or CHS study (available directly or as next best proxy in our meta-analysis), we estimated the “leave-two-out” (L2O) genetic effect β_L2O_ and standard error SE_L2O_ of the Holliday et al. meta-analysis as follows:

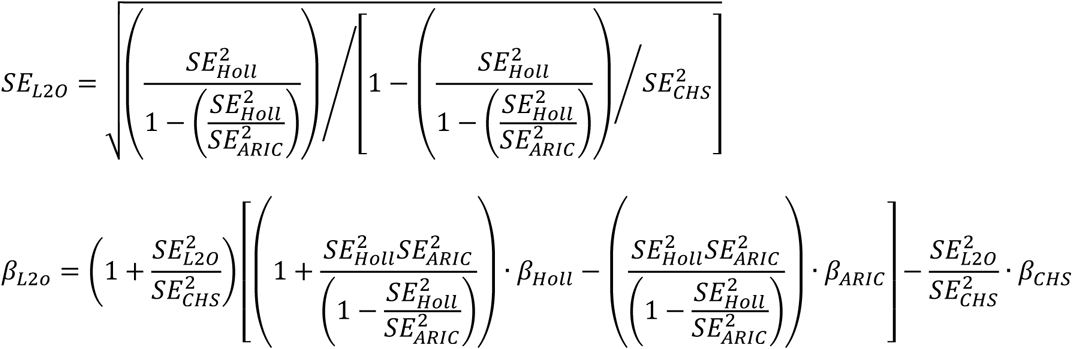

We applied a Wald test to test the corrected genetic effect for significance.

### Candidate approach

Additionally to the genome-wide search in our meta-analysis of 14,034 cases and 91,214 controls, we adopted a candidate approach based on the 14 reported suggestive variants by Holliday et al. (P-values from 8.9×10^-6^ to 1.1×10^-6^ in their meta-analysis, 4,089 cases and 20,453 controls). For this, we removed the overlapping studies from our meta-analysis (ARIC, CHS utilizing the reported variant; yielding 13,450 cases and 84,942 controls) and judged the variants’ association at a Bonferroni-corrected significant association (P< 0.05/14).

### Gene prioritization at newly identified AMD loci

To prioritize genes and variants at the newly identified AMD loci, we conducted a range of statistical and functional follow-up analyses. The following criteria were used: (1) Statistical evidence; we computed the Bayes Factor based posterior probability of each variant using Z-scores and derived 95% credible intervals for each locus (27). The method assumes a single causal signal per locus. (2) Variant effect predictor (VEP) to explore whether any of the credible variants was located in a relevant regulatory gene region (28). (3) eQTL analysis: We downloaded expression summary statistics for the candidate genes in retina from the EyeGEx database (29) and for 44 other tissues from the GTEx database (30) (both available at www.gtexportal.org/home/datasets) and evaluated whether any of the credible variants showed significant effects on expression levels in the aggregated data. For each significant eQTL in EyeGEx, we conducted colocalization analyses using eCAVIAR (31) to evaluate whether the observed early AMD association signal colocalized with the variants’ association with gene expression. (4) Retinal expression: We queried the EyeIntegration database to evaluate genes in the relevant loci for expression in fetal or adult retina or RPE cells (32). (5) Animal model: We queried the Mouse Genome Informatics (MGI) database (www.informatics.jax.org) for each gene in the relevant loci for relevant eye phenotypes in mice (33). (6) Human phenotype: The Online Mendelian Inheritance in Man (OMIM) ^®^ database was queried for human eye phenotypes (McKusick-Nathans Institute of Genetic Medicine, Johns Hopkins University, Baltimore, MD, queried 07/11/2019, www.omim.org).

### Phenome-wide association study for newly identified AMD loci

We used 82 other traits and queried reported genome-wide significant (P<5.0×10^-8^) lead variants and proxies (r^2^>0.5) for any of these traits for overlap with genes underneath our novel loci as done previously (34). For this, we used GWAS summary results that were previously aggregated from GWAS catalogue (35), GWAS central (36) and literature search.

For the novel early AMD lead variants, we further evaluated their association with 118 non-binary and 660 binary traits from the UK Biobank (37). The Phenome-wide association study (PheWAS) web browser “GeneATLAS” (www.geneatlas.roslin.ed.ac.uk) was used for the UK Biobank lookup. For each variant, association P values were corrected for the testing of multiple traits by the Benjamini-Hochberg false-discovery-rate (FDR) method (38).

### Interaction analyses

For the novel early AMD effects and for the 34 known advanced AMD lead variants (9), we investigated whether age modulated early AMD effects by analyzing variant x age interaction in seven data sources for which we had individual participant data available in Regensburg (ARIC, CHS, WHI, IAMDGC, UKBB, AugUR and KORA). For each source, we applied logistic regression and included a variant x AGE interaction term in the model (in addition to the covariates used in the main analysis). We conducted meta-analysis across the seven sources to obtain pooled variant x age interaction effects and applied a Wald test to test for significant interaction (at a Bonferroni-corrected alpha-level). For the novel early AMD effects, we further investigated whether age modulated advanced AMD effects by evaluating publically available data from IAMDGC (39). Finally, we investigated whether a novel early AMD lead variant modulated any of the effects of the 34 known AMD variants on advanced AMD (9). We used the IAMDGC data and applied one logistic regression model for each pair of known advanced AMD variants and novel early AMD variants including the two respective variants and their interaction (and the same other covariates as before).

### Comparison of genetic effects on early and advanced stage AMD

We estimated the genetic correlation between early and advanced AMD by utilizing the LDSC tool (40) with the GWAS summary statistics for early and advanced AMD (from the current meta-analysis and the IAMDGC (9), respectively). We used pre-calculated LD scores for European ancestry (https://data.broadinstitute.org/alkesgroup/LDSCORE/eur_w_ld_chr.tar.bz2). We further compared genetic effect sizes between early and advanced AMD for the novel early AMD lead variants and for the 34 known advanced AMD lead variants (9). For this, we queried the novel early AMD lead variants in the IAMDGC GWAS for advanced AMD (9) and (vice-versa) queried the 34 known advanced AMD lead variants (9) in the early AMD meta-analysis results. We compared effect sizes in a scatter plot and clustered the lead variants by their nominal significant association on advanced and/or early AMD. We classify different types of loci in a similar fashion as done previously for adiposity trait genetics (41): (1) “advanced-and-early” AMD loci (P_early_<0.05, P_adv_<0.05), (2) “advanced-only” AMD loci (P_early_≥0.05, P_adv_<0.05), (3) “early-only” AMD loci (P_early_<0.05, P_adv_≥0.05).

### Pathway analysis

To evaluate whether “advanced-and-early” AMD loci versus “advanced-only” AMD loci distinguished the major known pathways for advanced AMD, we performed pathway enrichment analysis separately for these two classes. We used the genes in the gene prioritization for all advanced AMD loci as previously described (9), derived the gene prioritization score and selected the best scored gene in each locus (two genes in the case of ties). We then separated the gene list according to the class of the respective locus, and performed pathway enrichment analysis via Enrichr (42) with default settings searching Reactome’s cell signaling pathway database 2016 (n=1,530 pathways). P-values were corrected for multiple testing with Benjamini-Hochberg procedure (38).

## RESULTS

### Eight genome-wide significant loci from a GWAS on early AMD

We conducted a meta-analysis of genotyped and imputed data from 11 sources (14,034 early AMD cases, 91,214 controls, for study-specific genotyping, analysis and quality control (**Tables S1-S2)**. For all participants, early AMD or control status (i.e. no early nor late AMD) was ascertained via color fundus photographs (**Table S1**). This included automated machine-learning based AMD classification of UK Biobank fundus images (application number 33999; 56,699 individuals from baseline, 13,650 additional individuals from follow-up) (15,19).

Based on logistic regression association analysis in each of the 11 data sets meta-analyzed via fixed effect model, we identified eight distinct loci with genome-wide significance (P=1.3×10^-116^ to 4.7×10^-8^, **Figure 1, Table 1**; “locus” defined by the lead variant and proxies, r^2^≥0.5, +/-500 kb). Six of these loci were novel for early AMD; two loci had been identified for early AMD previously (11).

**Table 1.**
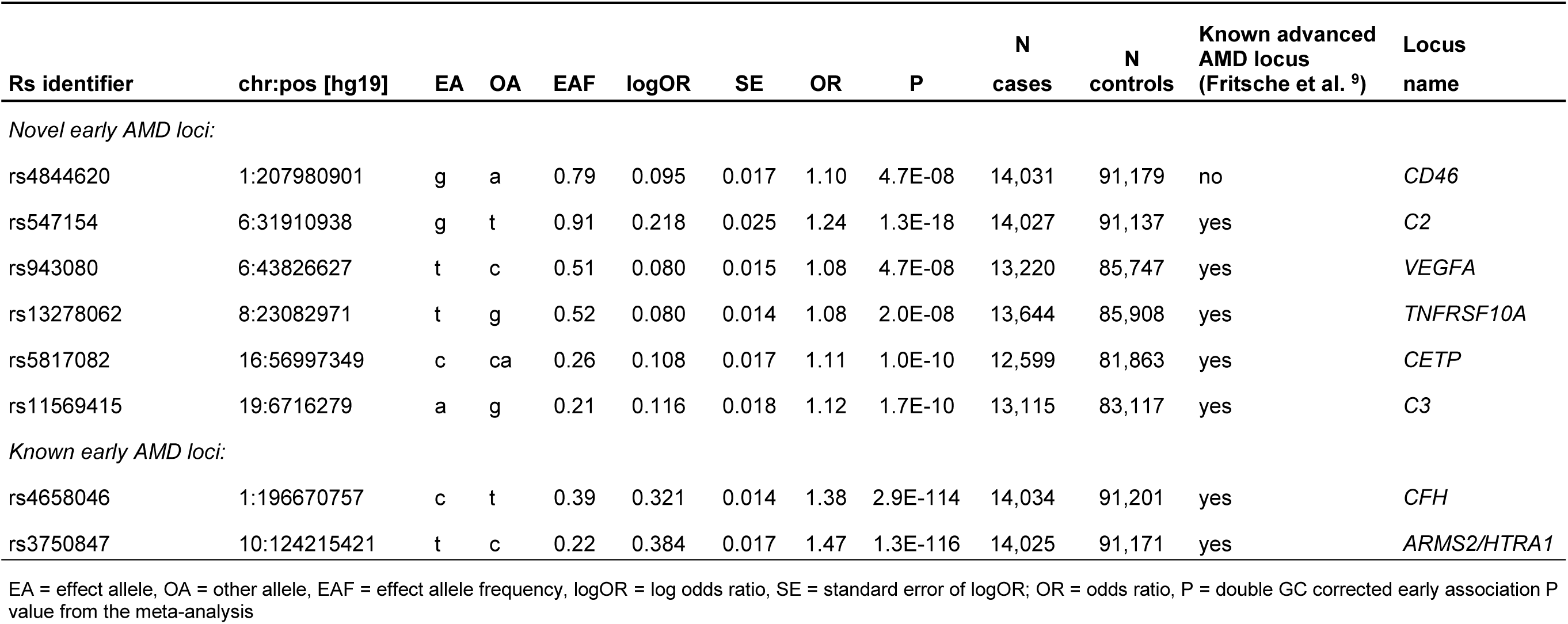
Genome-wide search for early AMD association. The table shows the eight genome-wide significant (P<5×10^-8^) lead variants from the early AMD meta-analysis.

**Fig 1.**
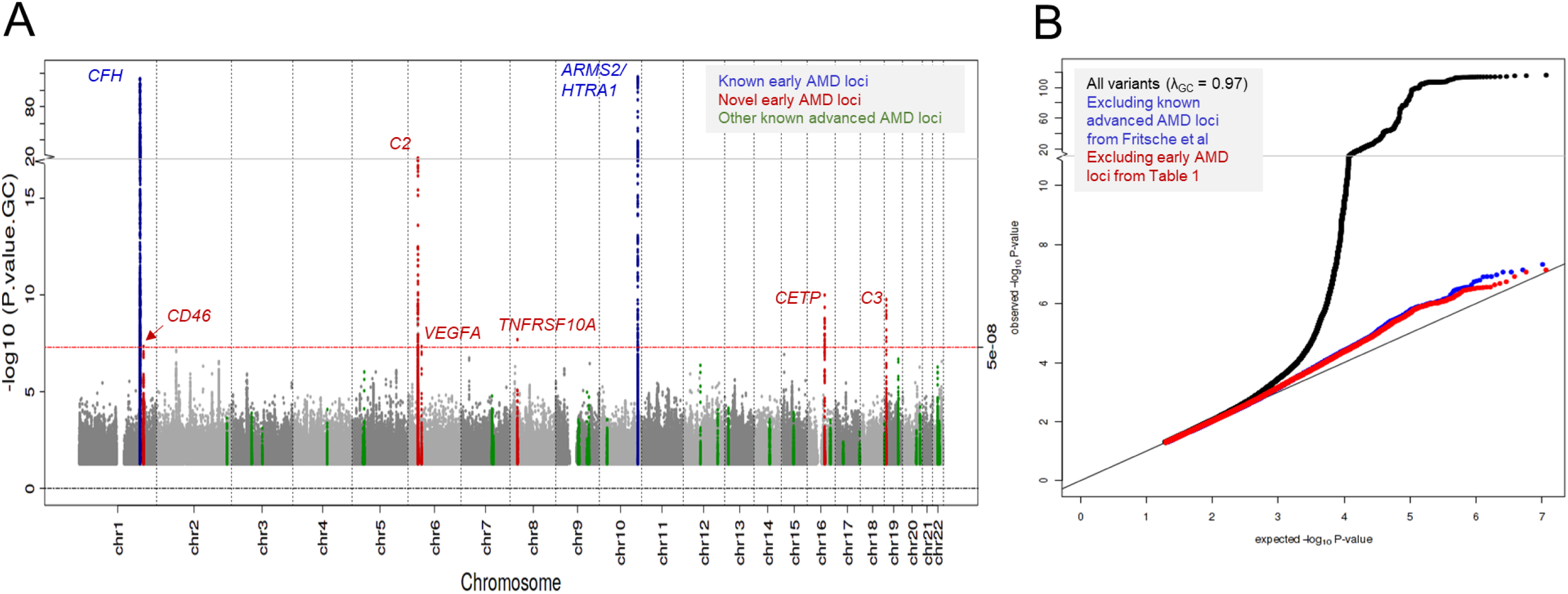
Early AMD meta-analysis. Shown are the association P-values of the meta-analysis for early AMD by their position on the genome (A, Manhattan plot) as well as their distribution (B, QQ plot). In A, color indicates whether the locus was previously identified by Holliday et al (11) (blue), novel for early AMD (red), or among the other advanced AMD loci identified by Fritsche et al (9) (green).

Most of these loci overlap with known loci for advanced AMD (9) (*CFH, ARMS2*/*HTRA1, C2, C3, CETP, VEGFA, TNFRSF10A)*, except one which has not been identified in early or advanced AMD GWAS before (P=4,7×10^-8^, lead variant rs4844620, near *CD46*, **Figure S1**). This novel locus is additionally supported by independent data from Holliday et al. (11) (3,432 cases and 11,235 controls after removing the overlap to our meta-analysis: P=7.0×10^-5^).

This novel locus on chromosome 1 is >10 million base positions distant from the closest known advanced AMD locus (around *CFH*) and the lead variant is uncorrelated with any of the eight independent *CFH* locus variants known for advanced AMD (r^2^<0.01). This locus showed no second signal (GCTA (26) conditional P≥5.0×10^-8^, **Figure S1**). Since early AMD ascertainment and classification was heterogeneous across the 11 data sources of our meta-analysis, we conducted sensitivity analysis leaving out one data set at a time and found the *CD46* locus association to be robust, except for a slightly larger early AMD effect when excluding IAMDGC data (**Figure S2**). Taken together, we identified six novel loci for early AMD including a novel locus for any AMD with genome-wide significance near *CD46*.

### Three significant loci from a candidate-based approach of 14 variants

Subsequently, we applied a candidate-based approach by investigating the lead variants of the 13 loci reported as suggestive by the previous GWAS for early AMD (4,089 early AMD cases, 20,453 controls; reported P between 1.1×10^-6^ and 8.9×10^-6^) (11); one further variant reported as suggestive was in the *CD46* locus that we have identified with genome-wide significance in our data and for which we have already utilized parts of the reported Holliday et al. data for independent replication (see above). We re-analyzed our data excluding the overlap with the previous GWAS (i.e. excluding ARIC, CHS study, yielding 13,450 early AMD cases and 84,942 controls). We found three of the 13 variants as significantly associated at a Bonferroni-corrected level (P<0.05/13=0.0038) depicting three novel loci for early AMD (**Table 2**). This included two variants in known advanced AMD loci (*APOE/TOMMS; PVRL2*) and one completely novel for any AMD near *TYR* (rs621313, P=6.8×10^-4^, **Figure S3**). There was no second signal within this locus (conditional P>0.0036, **Figure S3**). The observed effect was robust upon exclusion of any of the 11 data sets (**Figure S4**). Altogether, we have thus identified 9 novel loci for early AMD (6 from GWAS, 3 from candidate-based approach) including two completely novel for any AMD (1 from GWAS near *CD46*, 1 from candidate-based approach near *TYR*).

**Table 2.** Candidate approach to search for early AMD association. The table shows results for the 13 lead variants reported as suggestive for early AMD by Holliday et al. (11) (effective sample size = 13,631) for their association with early AMD in our data set (P<0.05/13= 0.0038, tested at Bonferroni-corrected significance level, effective sample size up to 42,119), excluding the variant in the *CD46* locus that we have already identified with genome-wide significance (see Table 1). The ARIC and CHS studies were excluded from our meta-analysis data to avoid overlap with the data by Holliday et al. (11).

| Rs identifier | chr:pos [hg19] | Locus<br>(Holliday et al. <sup>11</sup>) | Holliday et al. (11) |  |  |  |  | this meta-analysis (excluding ARIC, CHS) |  |  |  | Known advanced<br>AMD locus<br>(Fritsche et al. <sup>9</sup>) |
| --- | --- | --- | --- | --- | --- | --- | --- | --- | --- | --- | --- | --- |
|  |  |  | EA | OA | EAF | OR [CI 95%] | P | EAF | OR [CI 95%] | P | Neff |  |
| Novel loci for early AMD (P < 0.05/13) |  |  |  |  |  |  |  |  |  |  |  |  |
| rs621313 | 11:88913663 | TYR | A | G | 0.51 | 0.87 [0.83;0.92] | 3.5E-06 | 0.52 | 0.95 [0.93;0.98] | 6.8E-04 | 41,661 | no |
| rs6857 | 19:45392254 | PVRL2 | T | C | 0.15 | 0.81 [0.74;0.88] | 1.4E-06 | 0.16 | 0.92 [0.89;0.96] | 7.1E-05 | 38,938 | yes |
| rs2075650 | 19:45395619 | APOE/TOMM40 | A | G | 0.86 | 1.23 [1.13;1.34] | 1.1E-06 | 0.86 | 1.08 [1.04;1.13] | 2.6E-04 | 40,405 | yes |
| No significant association (P ≥ 0.05/13): |  |  |  |  |  |  |  |  |  |  |  |  |
| rs16851585 | 1:177568799 | - | C | G | 0.92 | 0.77 [0.69;0.86] | 5.0E-06 | 0.89 | 1.04 [0.99;1.09] | 0.099 | 42,119 | no |
| rs6721654 | 2:121301911 | GLI2,INHBB | T | C | 0.08 | 1.26 [1.14;1.4] | 6.5E-06 | 0.08 | 1.01 [0.96;1.07] | 0.65 | 41,718 | no |
| rs17586843 | 4:116924184 | - | T | C | 0.78 | 1.18 [1.1;1.27] | 1.5E-06 | 0.78 | 1.02 [0.98;1.05] | 0.31 | 42,119 | no |
| rs7750345 | 6:106260128 | - | A | G | 0.75 | 1.16 [1.09;1.24] | 6.8E-06 | 0.74 | 1.02 [0.98;1.05] | 0.32 | 42,119 | no |
| rs2049622 | 7:42176282 | GLI3 | A | G | 0.49 | 0.87 [0.83;0.93] | 8.9E-06 | 0.52 | 0.99 [0.96;1.02] | 0.41 | 42,060 | no |
| rs11986011 | 8:127332657 | FAM84B | T | C | 0.02 | 2.5 [1.68;3.71] | 5.0E-06 | - | - | - | - | no |
| rs6480975 | 10:54574996 | MBL2 | C | G | 0.84 | 1.21 [1.12;1.32] | 2.8E-06 | 0.85 | 0.99 [0.95;1.03] | 0.63 | 40,651 | no |
| rs4293143 | 11:82821382 | PCF11,RAB30 | T | G | 0.69 | 0.85 [0.79;0.91] | 7.8E-06 | 0.70 | 0.99 [0.96;1.02] | 0.50 | 42,119 | no |
| rs9646096 | 13:38065446 | POSTN,TRPC4 | A | C | 0.95 | 0.74 [0.65;0.84] | 6.0E-06 | 0.96 | 0.99 [0.92;1.06] | 0.76 | 39,782 | no |
| rs10406174 | 19:3944240 | ITGB1BP3,DAPK3 | A | G | 0.11 | 1.24 [1.13;1.36] | 5.6E-06 | 0.11 | 1.00 [0.93;1.07] | 0.92 | 17,936 | yes |

EA = effect allele, OA = other allele, EAF = effect allele frequency, OR = odds ratio, CI = confidence interval, P = P value from Holliday et al or Double GC corrected early association P value from this meta-analysis, Neff =
effective sample size

### Gene prioritization at the two novel loci

To prioritize variants and genes at the *CD46* and *TYR* locus, we conducted *in silico* follow-up analyses for all variants and overlapping genes for each of these two loci (4,451 or 5,729 variants, 10 or 7 genes, respectively). We found several interesting aspects (**Table 3**): (1) When prioritizing variants according to their statistical evidence for being the driver variant by computing 95% credible sets of variants (27), we found 23 and 294 credible set variants for the *CD46* and *TYR* locus, respectively (**Table S3**). (2) Using the Variant Effect Predictor (28), we assessed overlap of credible set variants with functional regulatory regions and found variants influencing the transcript and/or the protein for four genes (**Table S4**): variants causing an alternative splice form for *CD46*, a nonsense-mediated mRNA decay (NMD) for *CR1L*, a missense variant for *TYR* (rs1042602, r^2^=0.56 to the lead variant rs621313), and NMD variants for *NOX4.* (3) We investigated credible set variants for being an expression quantitative trait locus (eQTL) for any of the 17 genes in retina (Eye Genotype Expression database, EyeGEx (29)) or in 44 other tissues (Genotype-Tissue Expression database, GTEx (30)). For the *CD46* locus, we observed significant association of the lead variant and additional 16 credible set variants on *CD46* expression in retina (FDR<5%, **Table S5**); the early AMD risk increasing alleles of all 17 variants were associated with elevated *CD46* expression. Importantly, we observed the expression signal to colocalize with the early AMD association signal using eCAVIAR (31) (3 variants with colocalization posterior probability CLPP>0.01, **Table S6, Figure S5-S6**). We also found credible variants to be associated with *CD46* expression in 15 other tissues from GTEx, including four brain tissues (FDR<0.05, **Table S7).** Among the credible set variants in the two loci, we found no further eQTL for any of the other genes. When extending beyond the credible set, we found one further *CD46* locus variant as eQTL for *CD55*, but without colocalization (**Table S6, Figure S5-S6**). These findings support the idea that the credible set captures the essential signal. (4) We queried the 17 genes overlapping the two loci for expression in eye tissue and cells in EyeIntegration summary data (43). We found five and three genes, respectively, expressed in adult retina and adult RPE cells (*CD46, PLXNA2, CR1, CD34, CD55*; *TYR, GRM5, NOX4*; **Figure S7-S8**). (5) When querying the 17 genes in the Mouse Genome Informatics, MGI (33) or Online Mendelian Inheritance in Man, OMIM^®^, database, for eye phenotypes in mice or humans, we identified relevant eye phenotypes for five genes in mice (*CD46, CR1, CR1L, PLXNA2; TYR*; **Table S8**) and for one gene in human (*TYR;* **Table S9**).

**Table 3.** Summary of *in silico* follow-up and gene prioritization score (GPS). The table summarizes statistical and functional evidence for 10 and seven candidate genes of the novel early AMD loci on chromosome 1 and chromosome 11, respectively. Detailed results on the individual statistical and functional analyses are shown in **Tables S3-S9**. For the GPS, the sum of cell entries for “annotation” and “biology” was computed per row.

| Locus | Candidate gene | Chr | Pos-Start | Pos-End | Number of variants in 95% credible set | GPS | Annotation for variants in 95% credible set |  |  |  |  | Biology of the gene |  |
| --- | --- | --- | --- | --- | --- | --- | --- | --- | --- | --- | --- | --- | --- |
|  |  |  |  |  |  |  | Protein Altering | NMD | Altered splicing | eQTL * | Expressed in Eye tissue § | MGI Mouse eye phenotype | OMIM Human eye phenotype |
| CD46 | CD46 | 1 | 207925382 | 207968861 | 11 | 4 | 0 | 0 | 1 | 1 | 1 | 1 | 0 |
| CD46 | CR1L | 1 | 207818457 | 207897036 | 1 | 1 | 0 | 0 | 0 | 0 | 0 | 1 | 0 |
| CD46 | PLXNA2 | 1 | 208195587 | 208417665 | 0 | 2 | 0 | 0 | 0 | 0 | 1 | 1 | 0 |
| CD46 | CR1 | 1 | 207669472 | 207815110 | 0 | 2 | 0 | 0 | 0 | 0 | 1 | 1 | 0 |
| CD46 | LOC148696 | 1 | 207991723 | 207995941 | 1 | 0 | 0 | 0 | 0 | NA | NA | 0 | 0 |
| CD46 | CD34 | 1 | 208059882 | 208084683 | 0 | 1 | 0 | 0 | 0 | 0 | 1 | 0 | 0 |
| CD46 | CD55 | 1 | 207494816 | 207534311 | 0 | 1 | 0 | 0 | 0 | 0 | 1 | 0 | 0 |
| CD46 | CR2 | 1 | 207627644 | 207663240 | 0 | 0 | 0 | 0 | 0 | 0 | 0 | 0 | 0 |
| CD46 | MIR29B2 | 1 | 207975787 | 207975868 | 0 | 0 | 0 | 0 | 0 | NA | NA | 0 | 0 |
| CD46 | MIR29C | 1 | 207975196 | 207975284 | 0 | 0 | 0 | 0 | 0 | NA | NA | 0 | 0 |
| TYR | TYR | 11 | 88911039 | 89028927 | 39 | 4 | 1 | 0 | 0 | 0 | 1 | 1 | 1 |
| TYR | NOX4 | 11 | 89057521 | 89322779 | 1 | 3 | 0 | 1 | 1 | 0 | 1 | 0 | 0 |
| TYR | GRM5 | 11 | 88237743 | 88796846 | 109 | 1 | 0 | 0 | 0 | 0 | 1 | 0 | 0 |
| TYR | FOLH1B | 11 | 89392464 | 89431886 | 0 | 0 | 0 | 0 | 0 | 0 | 0 | 0 | 0 |
| TYR | GRM5-AS1 | 11 | 88237743 | 88257222 | 0 | 0 | 0 | 0 | 0 | 0 | NA | 0 | 0 |
| TYR | TRIM49 | 11 | 89530822 | 89541743 | 0 | 0 | 0 | 0 | 0 | 0 | NA | 0 | 0 |
| TYR | TRIM77 | 11 | 89443466 | 89451040 | 0 | 0 | 0 | 0 | 0 | 0 | NA | 0 | 0 |

\* Variants in 95% credible set are a local expression quantitative trait locus for this gene in retina (EyeGEx) or any tissue included in the GTEx database (cis for genes in locus); § Expression in Eye Integration data; NMD = nonsense-mediated mRNA decay; OMIM = Online Mendelian Inheritance in Man (https://www.omim.org/); NA = data not available; The gene start and end positions were extracted from the hg19 gene range list from http://www.cog-genomics.org/plink/1.9/resources.

While it is debatable how to prioritize evidence for a gene’s probability to be causal, one approach is to count any of the following characteristics for each of the 17 genes (Gene Prioritization Score, GPS **Table 3**): any credible set variant is (i) protein-coding, (ii) involved in NMD, (iii) affecting splice function, (iv) an eQTL for this gene in retina (EyeGEx) or in any other tissue (GTEx), or/and the gene (v) is expressed in retina or RPE, (vi) linked to eye phenotype in mouse or (vii) human. This approach offered *CD46* and *TYR* as the highest scored gene in the respective locus (GPS=4 for each; **Table 3**).

### Phenome-wide association search for the two novel loci

Co-association of variants in the two novel loci for early AMD with other traits and diseases may provide insights into shared disease mechanisms. We queried different data sets on numerous phenotypes by a gene-based and by a locus-based view.

For the gene-based view, we focused on 82 traits and evaluated reported genome-wide significant (P<5.0×10^-8^) lead variants (and proxies, r^2^>0.5) for overlap with any of the 17 gene regions (**Table S10**). For the *CD46* locus, we found significant association corrected for multiple testing (false-discovery rate, FDR < 5%) for schizophrenia (in *CD46* and *CR1L*) and for Alzheimer’s disease (in *CR1*, **Table S11**). For the *TYR* locus, we found significant associations for eye color, skin pigmentation and skin cancer (in *GRM5* and *TYR*, **Tables S11**).

For the locus-based view, we conducted a phenome-wide association study (PheWAS): we evaluated whether the two lead variants were associated with any of the 778 traits in UK Biobank using GeneAtlas (n=452,264, age-adjusted estimates; **Table S12**) (37). For the *CD46* lead variant, we identified 27 significant trait associations (FDR<5%), including four with particularly strong evidence (P<5.0×10^-8^; white blood cell, neutrophil, monocyte count and plateletcrit); the early AMD risk increasing allele (G, frequency=79%) was associated consistently with increased blood cell counts. We did not find a significant association of the *CD46* lead variant with schizophrenia in UK Biobank (FDR>5%; Alzheimer’s disease not available). For the *TYR* lead variant (rs621313, G allele associated with increased early AMD risk, frequency=48%), we identified 20 significant trait associations including Melanoma (FDR<5%, G allele associated with increased Melanoma risk) and two with particularly strong evidence for skin color and ease of skin tanning (P<5.0×10^-8^, G allele associated with brighter skin color and increased ease of skin tanning).

### Advanced AMD association and interaction analyses for the two novel loci

Next, we investigated whether the early AMD loci *CD46* or *TYR* were associated with advanced AMD. We thus queried the two lead variants for early AMD (rs4844620 and rs621313, respectively) for their advanced AMD association in the IAMDGC data (**Table S13**). We observed nominally significant directionally consistent effects for advanced AMD (OR_adv_=1.05, 95% confidence interval, CI=[1.01,1.09] and 1.03 [1.00,1.07], P_adv_=0.02 and 0.05, respectively) that were slightly smaller compared to early AMD effects (OR_early_=1.10 [1.06,1.14] and 1.05 [1.02,1.08], P_early_=4.7×10^-8^ and 6.8×10^-4^).

When exploring variant x age interaction for early AMD (in a subset of our meta-analysis of 10,890 early AMD cases and 54,697 controls) or for advanced AMD (IAMDGC data (39)) for the two novel locus lead variants, we found no statistically significant interaction at a Bonferroni-corrected level for early or advanced AMD (P_GxAGE_>0.05/2=0.025, **Table S14-S15**).

We were interested in whether one of the two novel lead variants showed interaction with any of the 34 known advanced AMD variants for association with advanced AMD (IAMDGC data). We found no significant interaction (P_GxG_>0.05/34/2), **Table S16**), which suggests that the known advanced AMD effects are not modulated by the two novel early AMD variants.

### Dissecting advanced AMD genetics into shared and distinct genetics for early AMD

We were interested in whether we could learn about advanced AMD genetics from a joint view of advanced and early AMD genetic effects. First, when computing genetic correlation of advanced AMD genetics with early AMD genetics, we found a substantial correlation of 78.8% (based on LD-score regression). Second, we contrasted advanced AMD effect sizes (IAMDGC data (9)) with early AMD effect sizes (our meta-analysis,) for the 34 known advanced AMD lead variants (**Figure 2, Table S13**). We found two classes of variants: (1) 25 variants showed nominally significant effects on early AMD (P<0.05; “advanced-and-early-AMD loci”), all directionally consistent and all smaller for early vs. advanced AMD (OR_early_=1.04-1.47; OR_adv_ =1.10-2.81); (2) nine variants had no nominally significant effect on early AMD (P≥0.05; “advanced-only AMD loci”). We did not find any variant with early AMD effects into the opposite direction as the advanced AMD effects. Also, we did not find any variant-age interaction on early AMD (**Table S15**).

**Fig 2.**
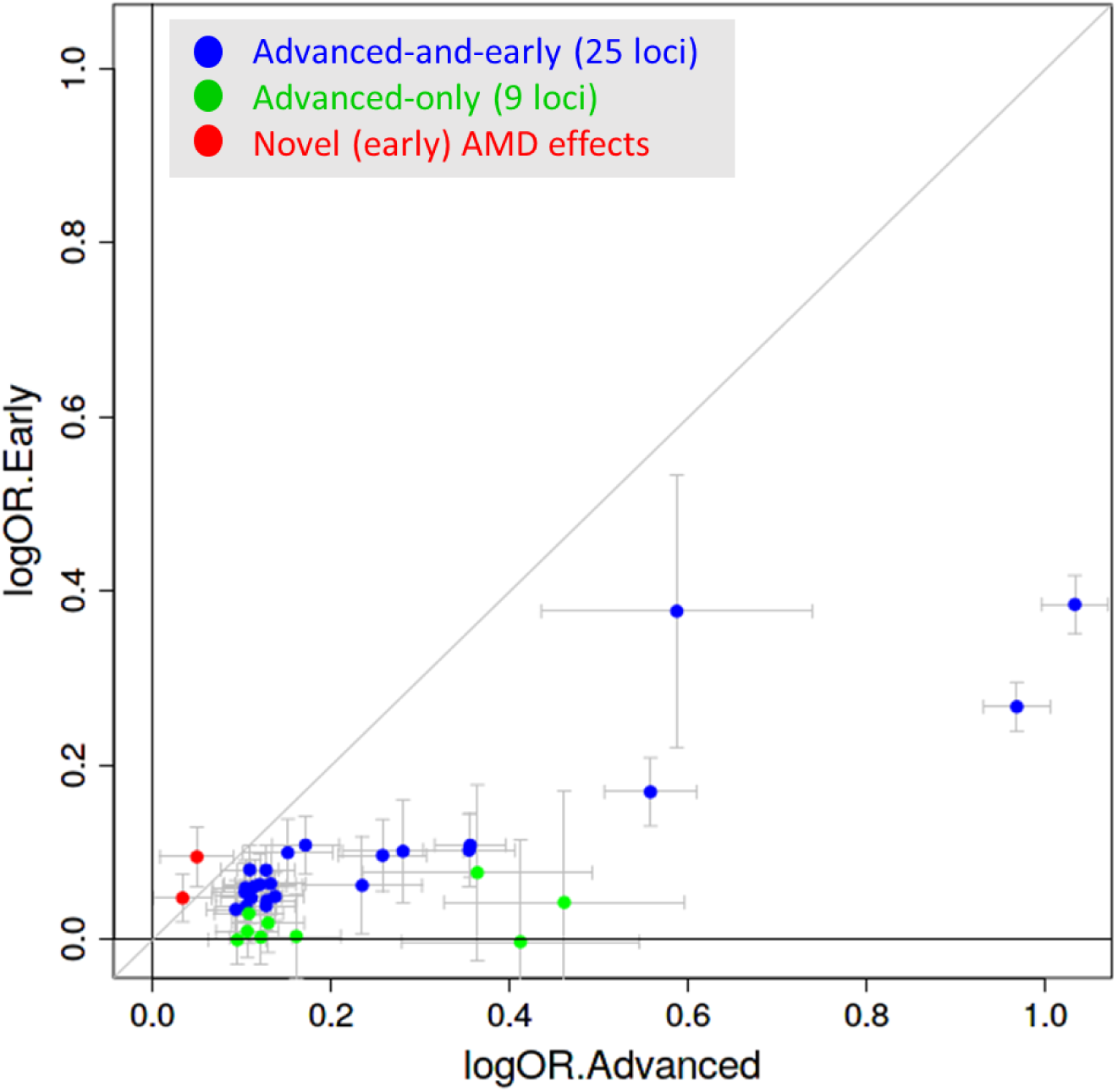
Advanced vs early AMD effect sizes. Shown are advanced AMD effect sizes contrasted to early AMD effect sizes (effect sizes as log odds ratios) for the 34 known advanced AMD variants (9) (blue or green for P_early_ < 0.05 or P_early_ ≥ 0.05, respectively) and for the two novel (early) AMD variants (red, near *CD46, TYR*). Detailed results are shown in **Table S13**.

We observed that complement genes *CFH, CFI, C3, C9*, and *C2* were all included in the 25 advanced-and-early-AMD loci. We were thus interested in whether advanced-and-early-AMD loci suggested different pathways compared to advanced-only-AMD loci. For this, we utilized the GPS from our previous work on advanced AMD (9) to select the best-supported genes in each of these loci (**Table S17**). We applied Reactome pathway analyses via Enrichr (42) twice: (i) for the 35 genes in the 25 advanced-and-early-AMD loci and (ii) for the nine genes in eight advanced-only-AMD loci (no gene in the “narrow” locus definition of the *RORB* locus). This revealed significant enrichment (corrected P<0.05) for genes from “complement system” and “lipoprotein metabolism” in the 25 advanced-and-early-AMD loci and enrichment for genes in the pathways “extracellular matrix organization” and “assembly of collagen fibrils” in the 8 advanced-only-AMD loci (**Table 4**). This suggested that the early AMD effect of advanced AMD variants distinguished the major known pathways for advanced AMD.

**Table 4.** Enriched pathways. The table shows enriched pathways for highest prioritized genes (from Fritsche et al. 2016 without modifications) in the 25 late AMD loci with early AMD effects (35 genes) versus the 8 loci with no effect on early AMD (9 genes). Pathways with significant corrected P-value (P_corr_ < 0.05) for each gene group from EnrichR querying human Reactome database 2016 are shown.

| Gene group | Reactome pathway | #Genes in gene set | #AMD loci in gene set | Raw P | Corrected P | Genes contributing to enrichment |
| --- | --- | --- | --- | --- | --- | --- |
| **Effects on early and advanced AMD** | Regulation of Complement cascade (R-HSA-977606) | 26 | 5 | $7.8 \times 10^{-10}$ | $1.2 \times 10^{-6}$ | *C3;CFH;C9;CFI;CFB* |
| | Lipoprotein metabolism (R-HSA-174824) | 34 | 4 | $3.5 \times 10^{-7}$ | $1.8 \times 10^{-4}$ | *ABCA1;CETP;LIPC;APOE* |
| | Complement cascade (R-HSA-166658) | 80 | 5 | $2.7 \times 10^{-7}$ | $2.0 \times 10^{-4}$ | *C3;CFH;C9;CFI;CFB* |
| | HDL-mediated lipid transport (R-HSA-194223) | 19 | 3 | $4.7 \times 10^{-6}$ | $1.8 \times 10^{-3}$ | *ABCA1;CETP;APOE* |
| | Lipid digestion, mobilization, and transport (R-HSA-73923) | 71 | 4 | $7.0 \times 10^{-6}$ | $2.2 \times 10^{-3}$ | *ABCA1;CETP;LIPC;APOE* |
| | Activation of C3 and C5 (R-HSA-174577) | 6 | 2 | $4.4 \times 10^{-5}$ | 0.011 | *C3;CFB* |
| **no effects on early AMD** | Assembly of collagen fibrils and other multimeric structures (R-HSA-2022090) | 54 | 3 | $1.5 \times 10^{-6}$ | $2.4 \times 10^{-3}$ | *COL15A1;COL8A1;MMP9* |
| | Collagen formation (R-HSA-1474290) | 85 | 3 | $6.1 \times 10^{-6}$ | $3.1 \times 10^{-3}$ | *COL15A1;COL8A1;MMP9* |
| | Extracellular matrix organization (R-HSA-1474244) | 283 | 4 | $4.7 \times 10^{-6}$ | $3.6 \times 10^{-3}$ | *VTN;COL15A1;COL8A1;MMP9* |

## DISCUSSION

Based on the largest genome-wide meta-analysis for early AMD to date encompassing ∼14,000 cases and ∼91,000 controls, all color fundus photography confirmed, we identified 11 loci for early AMD: 9 loci highlighted here for the first time with significant association for early AMD and two previously identified (11). Nine of the 11 loci overlapped with known loci for advanced AMD (9) and two had not been detected by GWAS for early or advanced AMD so far. Our post-GWAS approach highlighted *CD46* and *TYR* as compelling candidate genes in the two loci. Our joint view on early and advanced AMD genetics allowed us to differentiate between shared and distinct genetics for these two disease stages, which the pathway analyses suggested to be biologically relevant.

The locus around *CD46*, had not been identified with genome-wide significance by the previous largest GWAS for advanced AMD (9) (16,144 advanced AMD cases, 17,832 controls) or early AMD (11) (4,089 early AMD cases, 20,453 controls). Our meta-analysis was more than three times larger than the previous early AMD GWAS (effective sample size 48,651 compared to a 13,631(11)) and had a larger power to detect an “any AMD” effect with genome-wide significance than the previous advanced AMD GWAS (e.g. for OR=1.10, allele frequency 30%: power=92% compared to 61%, respectively). The *TYR* locus had not been genome-wide significant in any previous GWAS on advanced or early AMD; it was significant for early AMD at a Bonferroni-corrected level in our candidate approach (11).

Prioritization of genes underneath association signals is a known challenge, but highly relevant for selecting promising candidates for functional follow-up. Our systematic approach, scrutinizing all genes underneath our two newly identified loci, highlighted *CD46* and *TYR* as the most supported genes. *CD46* is an immediate compelling candidate as a part of the complement system (44). Complement activation in retina is thought to have a causal role for AMD (45,46). Importantly, we found our *CD46* GWAS signal to colocalize with *CD46* expression with the early AMD risk increasing allele (rs4844620 G) increasing CD46 expression in retinal cells. On the one hand, this contrasts the presumption that a higher *CD46* expression in eye tissue should protect from AMD, based on previous CD46 expression data (47) and a documented AMD risk increasing effect for increased complement inhibition (48). On the other hand, *CD46* had also been found to have pathogenic receptor properties for human viral and bacterial pathogens (e.g. measles virus) (49) and is known to down-modulate adaptive T helper type 1 cells (50). Furthermore, a GWAS on neutralizing antibody response to measles vaccine had identified two intronic *CD46* variants (rs2724384, rs2724374) (51). In our data, these two variants were in the 95% credible set for the *CD46* locus, highly correlated with our lead variant rs4844620 (r^2^>=0.95), and the major alleles (rs2724374 T, rs2724384 A) increased early AMD risk. Interestingly, the rs2724374 G was shown for *CD46* exon skipping resulting in a shorter CD46 isoform with a potential role in pathogen binding (51). Based on this, one may hypothesize that the observed *CD46* signal in early AMD is related to pathogenic receptor properties rather than complement inactivation.

At the second locus, *TYR* appears as the best supported gene by our systematic scoring. This locus and gene was already discussed by Holliday and colleagues (11), although they did not have statistically significant association with early AMD in their data. TYR is important for melanin production and *TYR* variants in human were associated with skin, eye and hair color (52–54). Melanin has a protective function in RPE against oxidative stress from UV radiation (55) and RPE pigmentation alterations are linked to AMD (56). In contrast to skin, melanin pigment in RPE was shown to be synthesized prenatally and stored in melanosomes throughout life, while tyrosinase activity in adult RPE remained controversial (57,58). Our study provides additional insight: our query of EyeIntegration data (32) found *TYR* expressed in adult human RPE, which may influence melanin production and its protective function in retina. Therefore, variants in or near *TYR* may represent risk factors beyond fetal melanin production. While we did not identify any *cis* effect between variants and *TYR* expression, one of our credible set variants in *TYR* (rs1042602) is a missense variant. Interestingly, this variant is a GWAS lead variant not only for skin color (53), but also for macular thickness in UK Biobank (59); the allele associated with thicker retina showed increased early AMD risk in our data. Since thicker RPE/Bruch’s membrane complex was associated with increased early AMD risk in the AugUR study (60), this would be in line with our early AMD risk increasing allele being linked to a process of increased accumulation of drusenoid debris in the RPE/Bruch’s membrane complex. Although *CD46* and *TYR* are the most supported genes in the two loci, we cannot rule-out the relevance of other genes in the loci.

It is a strength of the current study that early AMD and control status was ascertained by color fundus photography, not relying on health record data. However, the early AMD classification in our GWAS is heterogeneous across the 11 data sets: two studies incorporated information from optical coherence tomography (NICOLA, GHS), the UK Biobank classification was derived by a machine-learning algorithm (19), and the IAMDGC data was multi-site with different classification approaches (9). The uncertainty in early AMD classification and the substantial effort required for any manual AMD classification are likely reasons for the sparsity of early AMD GWAS so far. Our sensitivity analysis showed that our findings did not depend on one or the other data source or classification approach. Our data on early AMD genetics is comparable in size to the existing data on advanced AMD genetics from IAMDGC (summary statistics at http://amdgenetics.org/) and thus provides an important resource (summary statistics at http://genepi-regensburg.de) to enable a joint view.

By this joint view, we were able to differentiate the 34 loci known for advanced AMD into 25 “advanced-and-early-AMD loci” and nine “advanced-AMD-only loci”. Pathway enrichment analyses conducted separately for these two groups effectively discriminated the major known pathways for advanced AMD genetics (9): complement complex and lipid metabolism for “advanced-and-early-AMD” loci; extracellular matrix metabolism for “advanced-AMD-only” loci. The two novel loci around *CD46* and *TYR* fit to the definition of “advanced-and-early-AMD” loci and the *CD46* being part of the complement system supports the above stated pathway pattern. The larger effect size for early compared to advanced AMD for the two novel loci may – in part – be winner’s curse. How do our observations relate to potential etiological models? (1) For a genetic variant capturing an underlying mechanism that <u>triggers both early and advanced AMD</u>, we would expect the variant to show association with early <u>and</u> advanced AMD (compared to “healthy”) with directionally consistent effects (**Figure 3**, Model 1). This would be in line with our observed associations for the 27 “advanced-and-early-AMD” loci (25 known advanced AMD loci, 2 novel loci). This would also suggest that mechanisms of complement system or lipid metabolism trigger both early and advanced disease. (2) For a mechanism that <u>triggers advanced AMD</u> no matter whether the person is “healthy” or has early AMD, we would anticipate a variant effect for advanced AMD, but not for early AMD (**Figure 3**, Model 2). This would be in line with our observed associations for the nine “advanced-AMD-only” loci. This would also suggest that mechanisms of extracellular matrix metabolism trigger advanced AMD rather than early AMD. Of note, these include the *MMP9* locus, which is thought to trigger vascularization and wet AMD (9). (3) Another mechanism is conferred by variants that are purely responsible for <u>progression from early to advanced AMD</u>, but do not increase advanced AMD risk for “healthy” individuals. In such a scenario, the advanced AMD risk increasing allele would be under-represented among persons with early AMD (**Figure 3**, Model 3), particularly at older age, and it would be associated with decreased risk of early AMD (compared to “healthy). None of the identified variants showed this pattern overall or for older age in the variant x age interaction analyses. (4) For a mechanism that <u>triggers early AMD</u>, but has no impact on the progression from early to advanced AMD, we would have an effect on early AMD, but no effect on advanced AMD (**Figure 3**, Model 4). We did not find such a variant.

**Fig 3.**
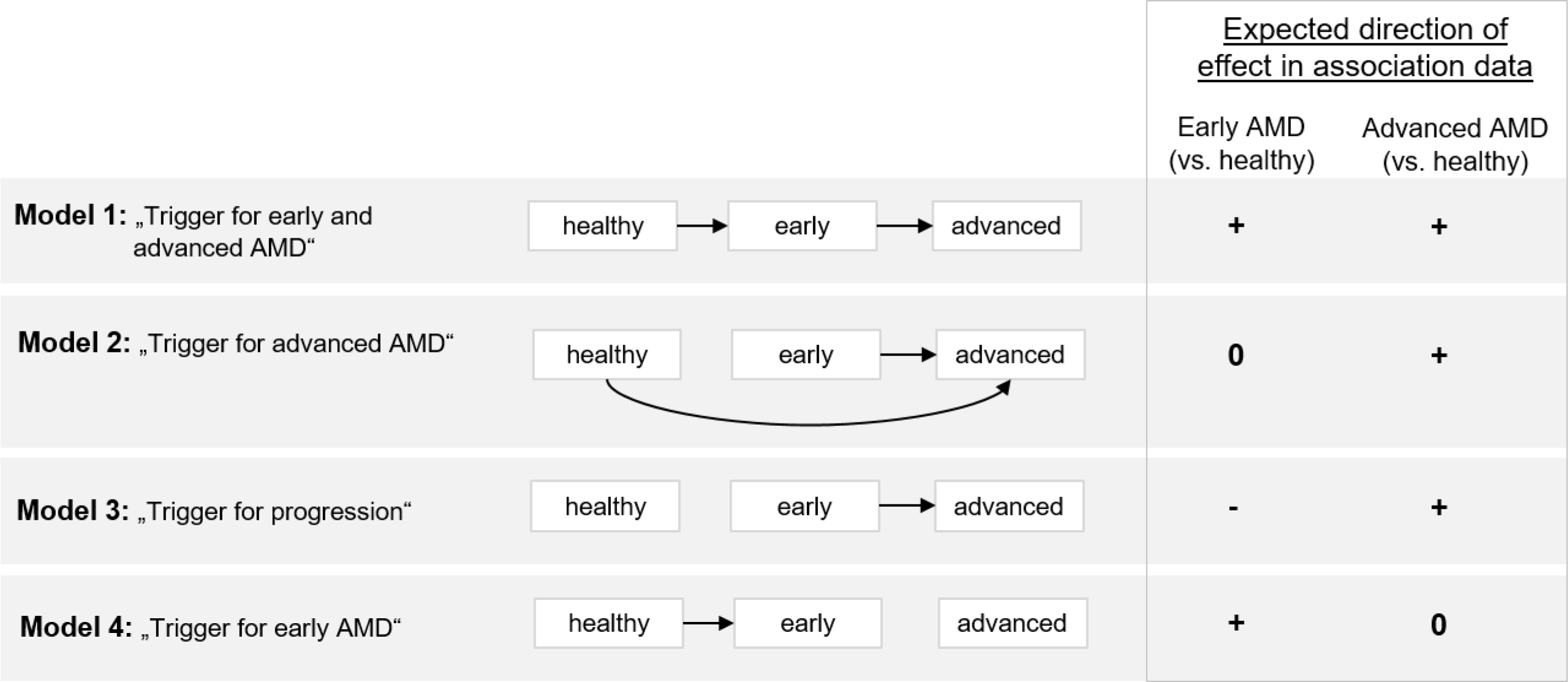
Etiological models and the respective expected association of a variant with early and advanced AMD.

Our data and joint view on effects for both disease stages support two of the four etiological models. One may hypothesize that the unsupported models are non-existing or unlikely. There are limitations to consider: (1) To reduce complexity, we adopted an isolated view per variant with some accounting for interaction, but ignoring more complex networks. (2) Early AMD effects were estimated predominantly in population-based studies, while advanced AMD effects were from a case-control design. (3) The cut-off of “nominal significance” for separating variants into “advanced-and-early” or “advanced-only” loci is arbitrary and larger data might give rise to re-classification. Still, the power to detect effects for early AMD in our meta-analysis was similar to the power in the advanced AMD data from IAMDGC (for OR=1.05, allele frequency 30%, nominal significance: power=94% or 83%, respectively). (4) An improved disentangling of genetic effects for the two chronologically linked disease stages will be an important subject of further research, requiring large-scale population-based studies with long-term follow-up and the estimation of transition probabilities.

## CONCLUSIONS

In summary, our large GWAS on early AMD identified novel loci, highlighted shared and distinct genetics between early and advanced AMD and provides insights into AMD etiology. The ability of early AMD effects to differentiate the major pathways for advanced AMD underscores the biological relevance of a joint view on early and advanced AMD genetics.

## LIST OF ABBREVIATIONS

3CC: Three Continent Consortium
AMD: Age-related macular degeneration
ARIC: The Atherosclerosis Risk in Communities Study
AugUR: Age-related diseases: Understanding Genetic and non-genetic influences - a study at the University of Regensburg
CHS: Cardiovascular Health Study eQTL: Expression quantitative trait locus FDR: False-discovery-rate
GCTA: Genome-wide Complex Trait Analysis
GC: Genomic control
GHS: Gutenberg Health Study
GPS: Gene Prioritization Score
GWAS: Genome-wide association study
HRC: Haplotype Reference Consortium
IAMDGC: International AMD Genomics Consortium
KORA: KOoperative Gesundheitsforschung in der Region Augsburg
LIFE: Leipzig Research Centre for Civilization Based Diseases - LIFE Adult population-based study, city of Leipzig, Germany
LOO: leave-one-out MAC: minor allele count
MGI: Mouse Genome Informatics
NICOLA: Northern Ireland Cohort for Longitudinal Study of Ageing NMD: nonsense-mediated mRNA decay
OCT: optical coherence tomography
OMIM: Online Mendelian Inheritance in Man
QC: quality control
RPE: retinal pigment epithelium
UKBB: UK Biobank
VEP: Variant effect predictor
WHI: Women’s Health Initiative

## DECLARATIONS

### Ethics approval and consent to participate

The Institutional Review Board (IRB) of the University of Utah was the umbrella IRB for all other studies contributing data to the International Age-related Macular Degeneration Genomics Consortium (IAMDGC), except for the Beaver Dam Eye Study (BDES). The University of Utah approved and certified each individual study ethic committee’s conduct for the data used in this study. Data provided by BDES was approved by the IRB of the University of Wisconsin. Local ethics approval for data access to the studies deposited in dbGAP (WHI, ARIC and CHS) was granted by the IRB of the University of Regensburg. For all other studies, study participants obtained informed consent and local ethics committees approved the study protocols.

### Consent for publication

Not applicable.

### Availability of data and materials

The genome-wide meta-analysis summary statistics are available for download from http://genepi-regensburg.de.

### Competing interests

M.S. receives funding from Pfizer Inc. for a project not related to this research. Retinal grading of the NICOLA study was supported by Novartis (for R.E.H) and Bayer (for Usha Chakravarthy, not a co-author). A.K.S. received financial and research support by Heidelberg Engineering, Novartis, Bayer Vital and Allergen without a link to the content of this work. I.M.H. received funding from Roche Diagnostics for a project not related to this research. None of the other authors have any conflicts of interest.

### Funding

The International AMD Genomics Consortium (IAMDGC) is supported by a grant from NIH (R01 EY022310). Genotyping was supported by a contract (HHSN268201200008I) to the Center for Inherited Disease Research (http://amdgenetics.org/). In-depth analyses to estimate genetic effects in the IAMDGC data was supported by DFG HE 3690/5-1 to I.M.H. The AugUR study was supported by grants from the German Federal Ministry of Education and Research (BMBF 01ER1206, BMBF 01ER1507 to I.M.H.) and the University of Regensburg. The KORA study was initiated and financed by the Helmholtz Zentrum München – German Research Center for Environmental Health, which is funded by the German Federal Ministry of Education and Research (BMBF) and by the State of Bavaria. Furthermore, KORA research was supported within the Munich Center of Health Sciences (MC-Health), Ludwig-Maximilians-Universität, as part of LMUinnovativ. This publication is supported by the Leipzig Research Centre for Civilization Diseases (LIFE), an organizational unit affiliated to the Medical Faculty of Leipzig University. LIFE is funded by means of the European Union, by the European Regional Development Fund (ERDF) and by funds of the Free State of Saxony within the framework of the excellence initiative (project numbers: 713-241202, 14505/2470, 14575/2470). Franziska G. Rauscher (F.G.R.) holds a grant from the German Federal Ministry of Education and Research: i:DSem - Integrative data semantics in systems medicine (031L0026). Tobias Elze (T.E.) is funded by the Lions Foundation, Grimshaw-Gudewicz Foundation, Research to Prevent Blindness, BrightFocus Foundation, Alice Adler Fellowship, NEI R21EY030142,NEI R21EY030631, NEI R01EY030575, and NEI Core Grant P30EYE003790. The Gutenberg Health Study is funded through the government of Rhineland-Palatinate („Stiftung Rheinland-Pfalz für Innovation”, contract AZ 961-386261/733), the research programs “Wissen schafft Zukunft” and “Center for Translational Vascular Biology (CTVB)” of the Johannes Gutenberg-University of Mainz, and its contract with Boehringer Ingelheim and PHILIPS Medical Systems, including an unrestricted grant for the Gutenberg Health Study. Alexander K Schuster (A.K.S.) holds the professorship for ophthalmic healthcare research endowed by „Stiftung Auge” and financed by „Deutsche Ophthalmologische Gesellschaft” and „Berufsverband der Augenärzte Deutschland e.V.”. The Atherosclerosis Risk in Communities study has been funded in whole or in part with Federal funds from the National Heart, Lung, and Blood Institute, National Institute of Health, Department of Health and Human Services, under contract numbers (HHSN268201700001I, HHSN268201700002I, HHSN268201700003I, HHSN268201700004I, and HHSN268201700005I). The research reported in this article was supported by contract numbers N01-HC-85079, N01-HC-85080, N01-HC-85081, N01-HC-85082, N01-HC-85083, N01-HC-85084, N01-HC-85085, N01-HC-85086, N01-HC-35129, N01 HC-15103, N01 HC-55222, N01-HC-75150, N01-HC-45133, N01-HC-85239 and HHSN268201200036C;grant numbers U01 HL080295 from the National Heart, Lung, and Blood Institute and R01 AG-023629 from the National Institute on Aging, with additional contribution from the National Institute of Neurological Disorders and Stroke. The WHI program is funded by the National Heart, Lung, and Blood Institute, National Institutes of Health, U.S. Department of Health and Human Services through contracts N01WH22110, 24152, 32100-2, 32105-6, 32108-9, 32111-13, 32115, 32118-32119, 32122, 42107-26, 42129-32, and 44221. This study has been conducted using the UK Biobank resource under Application Number 33999. The UK Biobank was established by the Wellcome Trust medical charity, Medical Research Council, Department of Health, Scottish Government and the Northwest Regional Development Agency. It has also had funding from the Welsh Assembly Government, British Heart Foundation and Diabetes UK. The analyses were supported by German Research Foundation (DFG HE-3690/5-1 to I.M.H.) and by the National Institutes of Health (NIH R01 EY RES 511967 to I.M.H.). Felix Grassmann (F.G.) was a Leopoldina Postdoctoral Fellow (Grant No. LPDS 2018-06) funded by the Academy of Sciences Leopoldina. The position of Tobias Strunz (T.S.) is financed by the Helmut-Ecker-Foundation (# 05/17 to B.H.F.W.) and of Christina Kiel (C.K.) by a grant from the German Research Foundation to F.G. and B.H.F.W. (GR 5065/1-1).

## Acknowledgments

Extended acknowledgments are shown in **Table S1**.

## Authors’ contributions

TWW, FGr, CB, CK, FGü, IMH, KJS and BHFW designed the study and wrote the manuscript. TWW and FGr conducted the meta-analysis. FGü applied the automated grading of UK Biobank fundus images. TWW and CK conducted the PheWAS. TWW, LW, MEZ and KJS conducted analysis for the gene priority scoring. KS conducted the pathway analyses. CAK, AP and AKS contributed data from the GHS study. MMN and AP contributed data from the KORA study. FGR, TE, KH, and MS contributed data from the LIFE-Adult study. MCG, AJMK, NQ and REH contributed data from the NICOLA study. IMH, KJS and BHFW supervised the study. All authors read and approved the final manuscript.

