## Supplementary Figures for "Genome-wide association meta-analysis for early age-related macular degeneration highlights novel loci and insights for advanced disease"

### SUPPORTING FIGURES

**Figure S1. CD46 regional association.** Shown are regional association plots for the *CD46* locus (A: primary meta-analysis, B: Approximate conditional GCTA analyses, conditioned on rs4844620). Color indicates the correlation (r²) to rs4844620.

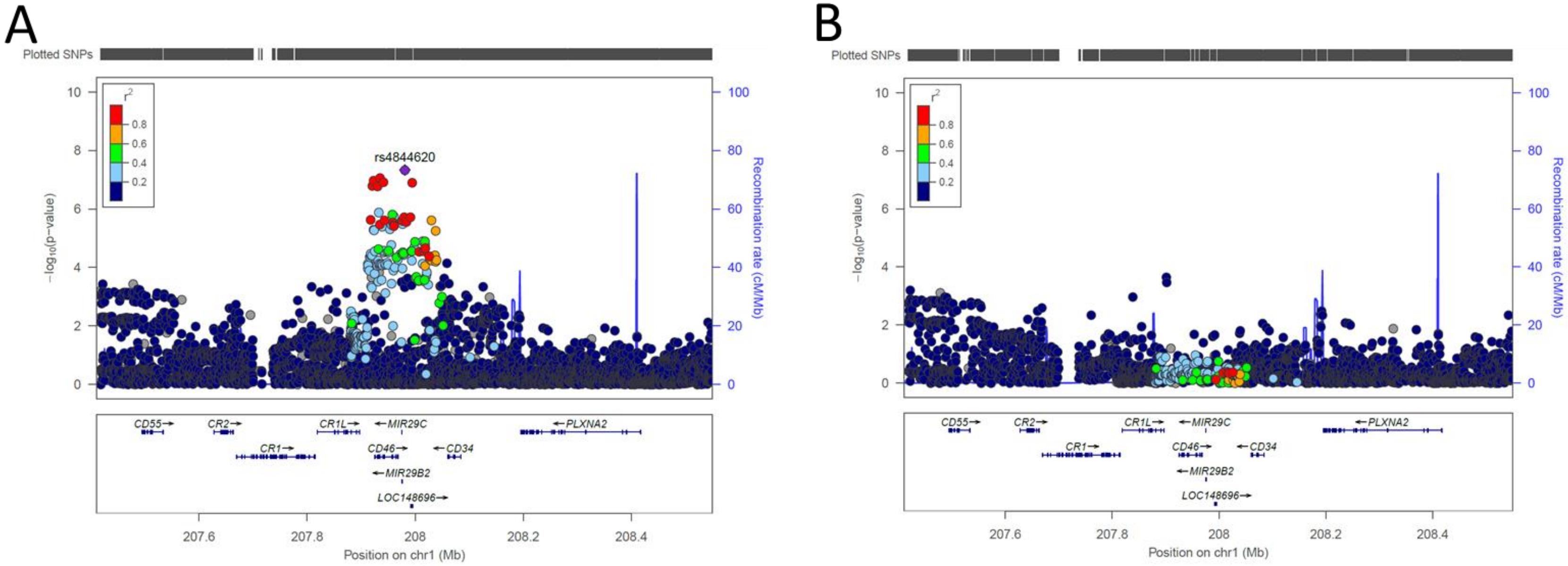

**Figure S2. Leave-one-out meta-analysis for the newly identified early AMD locus near *CD46*.** The forest plots show log odds ratios on early AMD with confidence intervals for rs4844620 (near *CD46*) from each of the 11 leave-one-study-out meta-analyses. The first column of each figure states the excluded study of the respective meta-analysis.

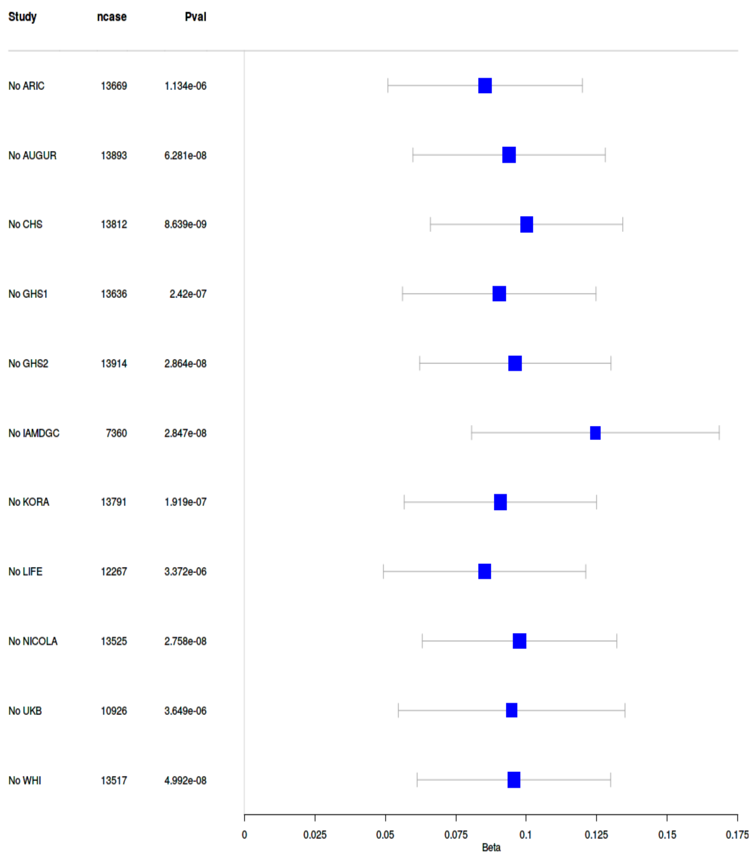

**Figure S3. TYR regional association.** Shown are regional association plots for the *TYR* locus (A: primary meta-analysis, B: Approximate conditional GCTA analyses, conditioned on rs621313). Color indicates the correlation (r²) to rs621313.

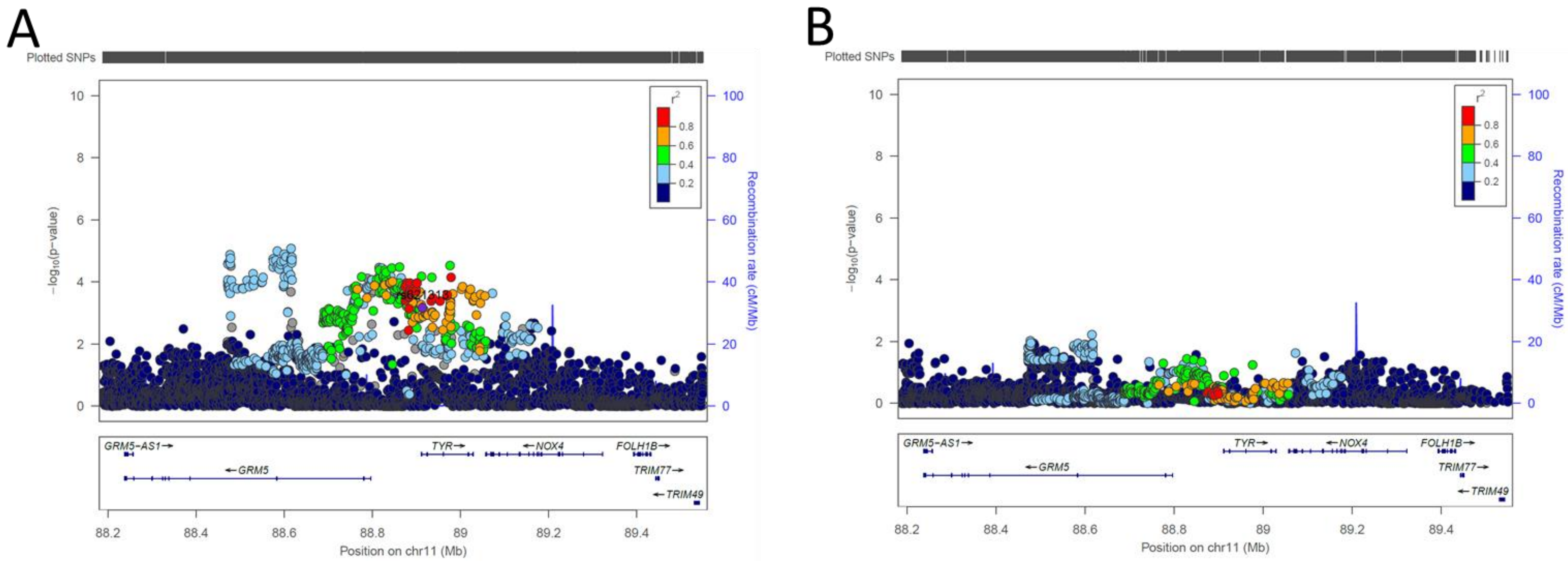

**Figure S4. Leave-one-out meta-analysis for the newly identified early AMD locus near *TYR*.** The forest plots show log odds ratios on early AMD with confidence intervals for rs621313 (near *TYR*) from each of the 11 leave-one-study-out meta-analyses. The first column of each figure states the excluded study of the respective meta-analysis.

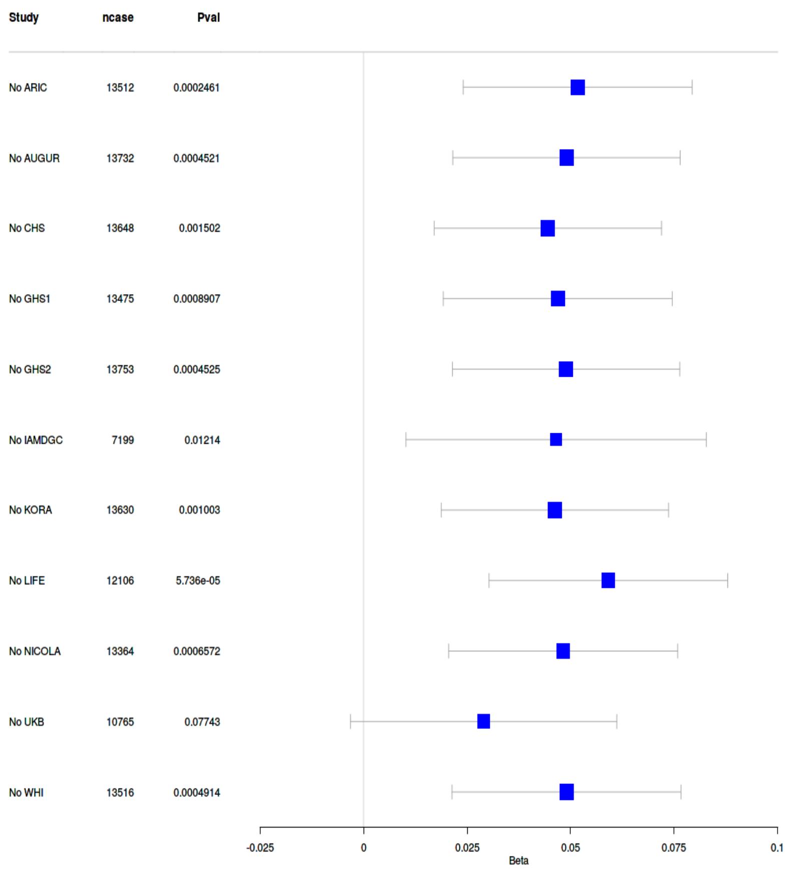

**Figure S5.** Colocalization of early AMD association and expression of *CD46* or *CD55* in human retina. The scatter plots compare association P values for early AMD (x axis) with association on expression of CD46 (panel A, y axis) or CD55 (panel B, y axis). The variants are colored by $r^2$ to the *CD46* lead variant rs4844620.

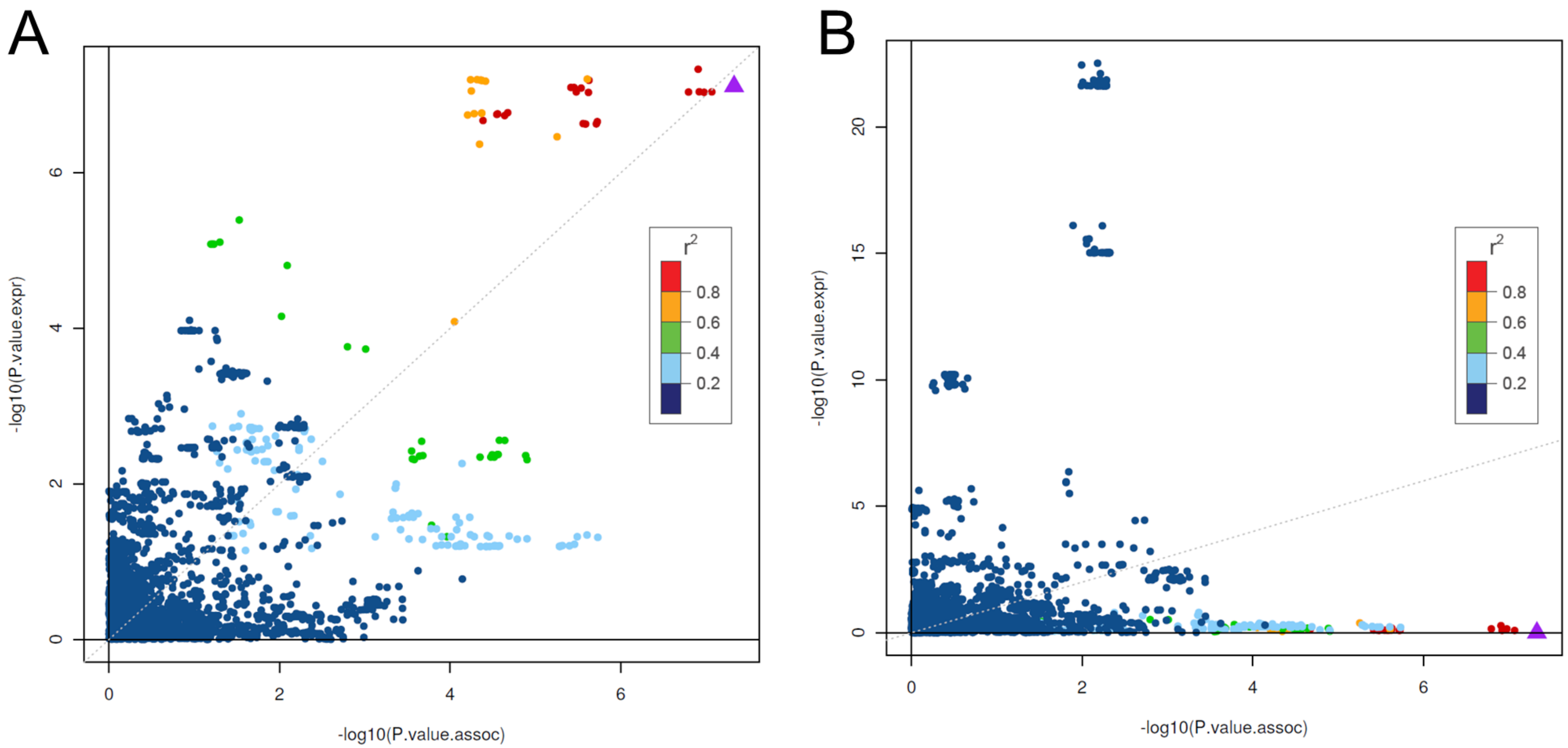

**Figure S6.** Colocalization of association on early AMD (meta-analysis) and expression of *CD46, CD34, PLXNA2* and *CD55* in human retina. The plots compare the association P values for expression (red) with early AMD (blue) by chromosomal base position (x axis).

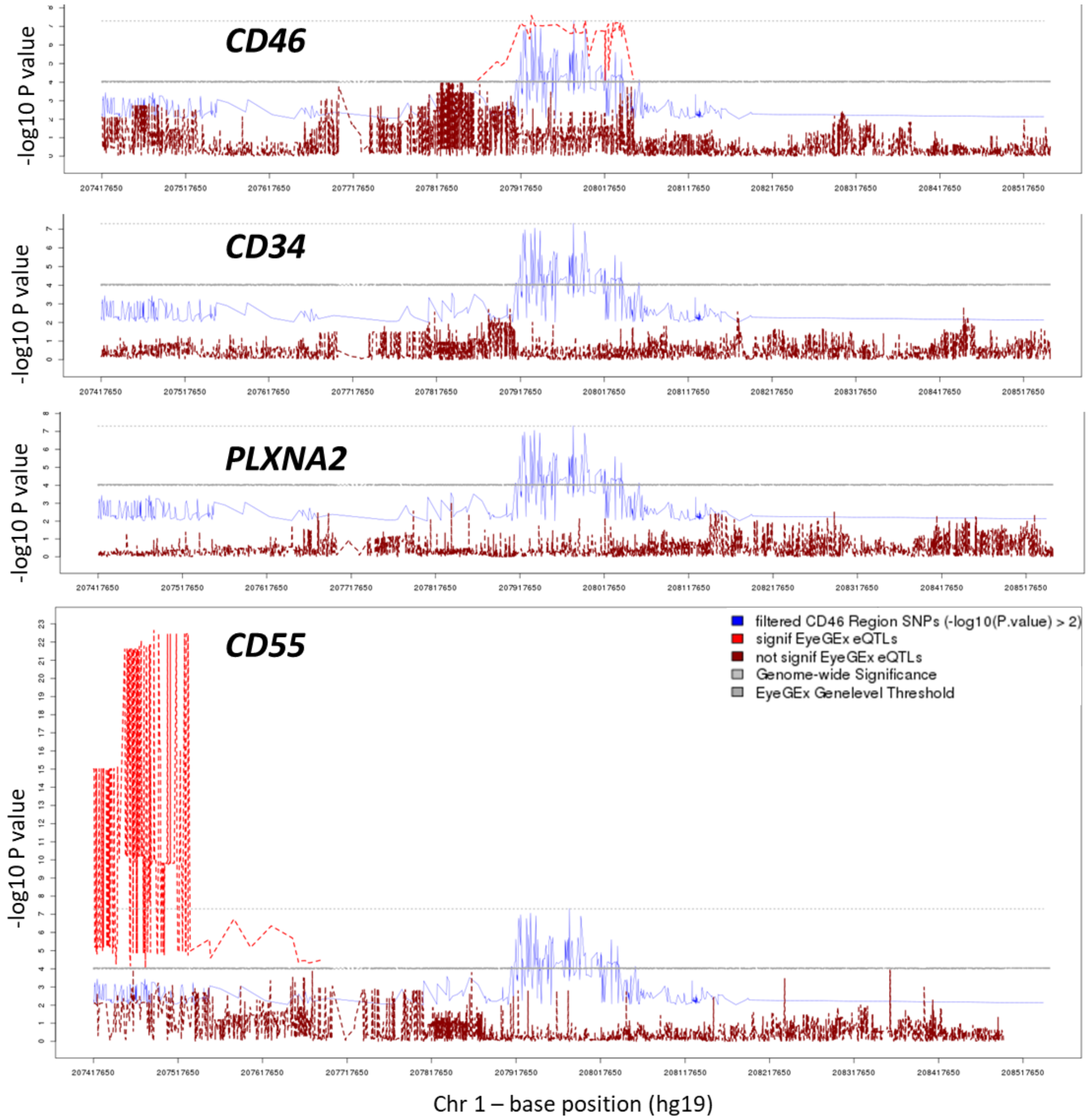

**Figure S7.** Expression of candidate genes at the *CD46* locus in EyeIntegration data for Fetal Retina (A), Adult Retina (B), Fetal RPE (C) and Adult RPE (D). The percentiles denote the distribution of gene expression across all genes (approximately 36,852 genes measured) in the respective tissue.

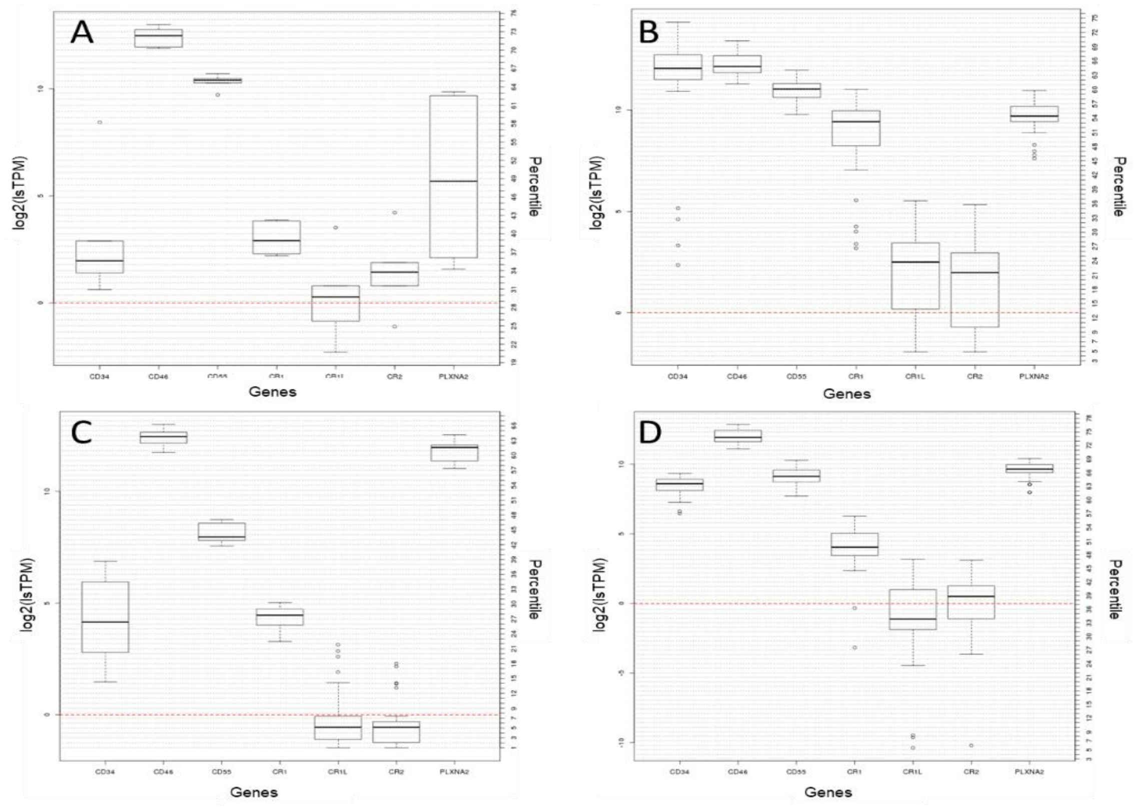

**Figure S8.** Expression of candidate genes at the *TYR* locus in EyeIntegration data for Fetal Retina (A), Adult Retina (B), Fetal RPE (C) and Adult RPE (D). The percentiles denote the distribution of gene expression across all genes (approximately 36,852 genes measured) in the respective tissue.

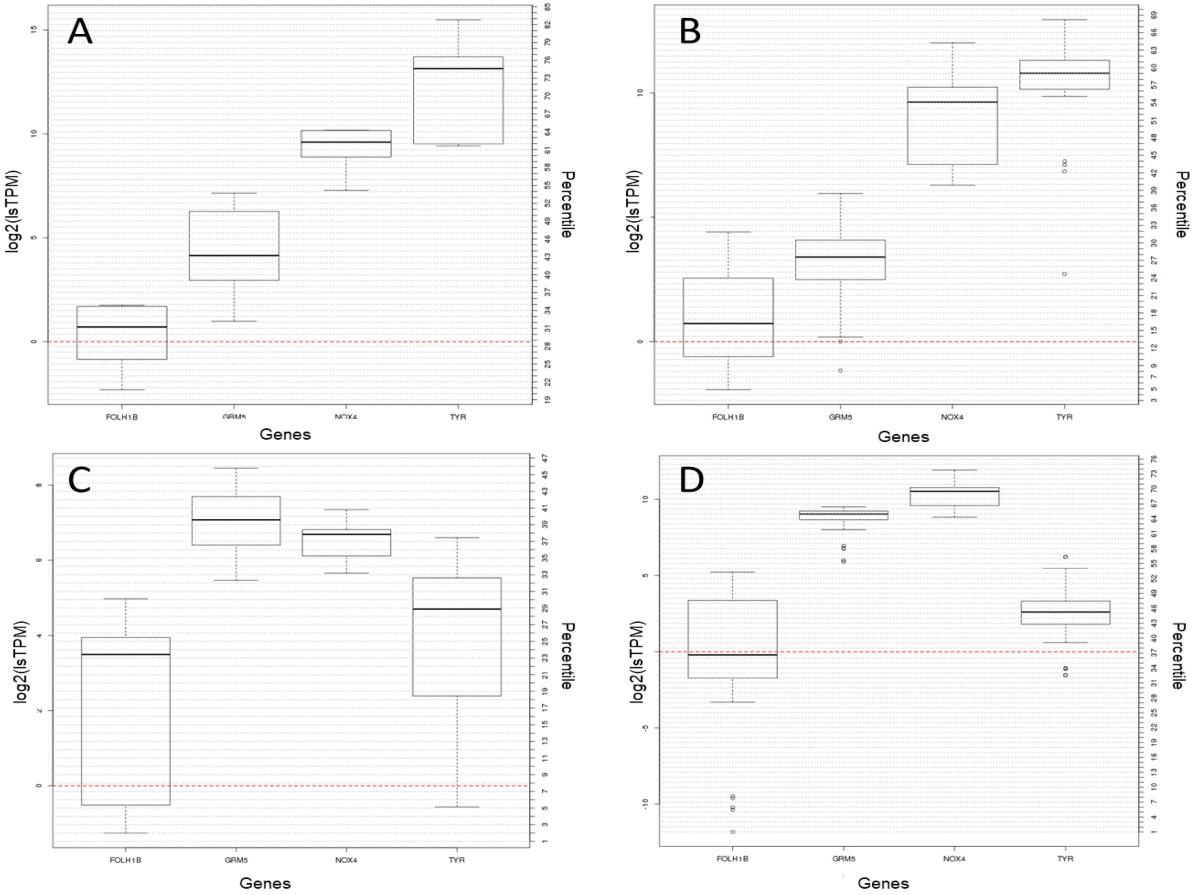
